# Degron-based bioPROTACs for controlling signaling in CAR T cells

**DOI:** 10.1101/2024.02.16.580396

**Authors:** Matthew S. Kim, Hersh K. Bhargava, Gavin E. Shavey, Wendell A. Lim, Hana El-Samad, Andrew H. Ng

**Affiliations:** Tetrad Graduate Program, University of California, San Francisco, San Francisco, CA; Cell Design Institute, University of California, San Francisco, San Francisco, CA; Biophysics Graduate Program, University of California, San Francisco, San Francisco, CA; Cell Design Institute, University of California, San Francisco, San Francisco, CA; Department of Biochemistry and Biophysics, University of California, San Francisco, San Francisco, CA; Department of Cellular and Molecular Pharmacology, University of California, San Francisco, San Francisco, CA; Arsenal Biociences, Inc., South San Francisco, CA; Cell Design Institute, University of California, San Francisco, San Francisco, CA; Cell Design Institute, University of California, San Francisco, San Francisco, CA; Department of Cellular and Molecular Pharmacology, University of California, San Francisco, San Francisco, CA; Altos Labs, Redwood City, CA; Cell Design Institute, University of California, San Francisco, San Francisco, CA; Department of Biochemistry and Biophysics, University of California, San Francisco, San Francisco, CA; Chan-Zuckerberg Biohub, San Francisco, CA; Department of Biochemistry and Biophysics, University of California, San Francisco, San Francisco, CA; Department of Molecular Biology, Genentech Inc., South San Francisco, CA, USA; Cell Design Institute, University of California, San Francisco, San Francisco, CA; Department of Cellular and Molecular Pharmacology, University of California, San Francisco, San Francisco, CA

## Abstract

Chimeric antigen receptor (CAR) T cells have made a tremendous impact in the clinic, but potent signaling through the CAR can be detrimental to treatment safety and efficacy. The use of protein degradation to control CAR signaling can address these issues in pre-clinical models. Existing strategies for regulating CAR stability rely on small molecules to induce systemic degradation. In contrast to small molecule regulation, genetic circuits offer a more precise method to control CAR signaling in an autonomous, cell-by-cell fashion. Here, we describe a programmable protein degradation tool that adopts the framework of bioPROTACs, heterobifunctional proteins that are composed of a target recognition domain fused to a domain that recruits the endogenous ubiquitin proteasome system. We develop novel bioPROTACs that utilize a compact four residue degron and demonstrate degradation of cytosolic and membrane protein targets using either a nanobody or synthetic leucine zipper as a protein binder. Our bioPROTACs exhibit potent degradation of CARs and can inhibit CAR signaling in primary human T cells. We demonstrate the utility of our bioPROTACs by constructing a genetic circuit to degrade the tyrosine kinase ZAP70 in response to recognition of a specific membrane-bound antigen. This circuit is able to disrupt CAR T cell signaling only in the presence of a specific cell population. These results suggest that bioPROTACs are a powerful tool for expanding the cell engineering toolbox for CAR T cells.

## Introduction

Engineered cells can sense environmental cues, process these cues through genetic circuits, and respond with therapeutic action. The potential of engineered cell therapies is highlighted by the clinical success of chimeric antigen receptor (CAR) T cells. CARs are composed of a custom extracellular domain, typically a single chain variable fragment (scFv), and signaling domains taken from the T cell receptor and associated co-stimulatory receptors. These synthetic receptors can redirect T cell signaling towards cancer cells, demonstrating complete response in over 85% of patients with blood cancers that were resistant to other therapeutics such as chemotherapy^1,2^. However, potent and sustained CAR signaling can be a double-edged sword. High levels of CAR T cell activity are associated with cytokine release syndrome (‘CRS’) resulting in severe inflammation, life-threatening shock and organ failure. Furthermore, chronic activation of the CAR can induce a hypofunctional state known as T cell exhaustion, limiting the therapeutic efficacy of CAR T cell therapies^3,4^. Regulating CAR signaling in T cells has the potential to improve therapeutic efficacy and safety.

Targeted protein degradation (TPD) is an exciting method of modulating cell signaling^5^. Work in preclinical models has shown that small molecule-induced degradation of the CAR protein can inhibit signaling and therapeutic activity, and mitigate CAR T cell exhaustion^6,7,8^ While these small molecule-based systems are capable of reversible and potent degradation, they systemically downregulate CAR expression. This non-specific degradation could hamper global CAR T cell therapeutic efficacy.

Regulating CAR expression via genetic circuits enables cell-by-cell decision making to modulate T cell signaling in response to local environmental cues, enhancing therapeutic efficacy and safety. A key example of genetic circuits in cellular therapies are AND logic gates that utilize antigen-sensing receptors such as synthetic Notch (synNotch) and Synthetic Intramembrane Proteolysis Receptors (SNIPR) to activate CAR expression. SynNotch and SNIPRs are force-sensitive chimeric receptors composed of an extracellular antigen binding domain, transmembrane and juxtamembrane domains from mammalian Notch receptors and an intracellular synthetic transcription factor. Antigen binding triggers proteolytic cleavage and release of the transcription factor from the plasma membrane to trigger expression of a custom genetic payload. This AND gate circuit topology improves CAR T cell specificity and safety in mouse models and has even been shown to alleviate exhaustion^9–14^. Genetic circuits have also been constructed with cell-state promoters or synNotch to drive cytokine production in response to specific cellular contexts to improve CAR T cell tumor clearance^15,16^. These examples demonstrate the potential for genetic circuits to enhance engineered cell therapies.

The coupling of genetic circuits with protein degradation could give rise to improved strategies for controlling CAR T cell signaling. However, existing small molecule-based protein degradation methods for CAR control are not amenable to incorporation into genetic circuits. An alternative method of inducing degradation of a protein of interest (POI) is the use of protein-based heterobifunctional molecules known as bioPROTACs. bioPROTACs are modular molecules composed of one protein domain that binds a POI and another protein domain that promotes ubiquitination of the POI^17–21^. Endogenous protein binding domains or motifs and engineered proteins such as DARPins, nanobodies and monobodies, have been successfully used to target a wide variety of proteins, including HER2, PCNA,and KRAS, for degradation using bioPROTAC molecules^22–27^. For the majority of bioPROTACs, truncated endogenous E3 ligases have been utilized to promote ubiquitination of a POI^28^. Interestingly, degrons, short degradation inducing sequences, can replace these much bulkier protein domains while retaining degradation activity^18,29^. Degrons known to interact with E3 ligases have been screened for development of bioPROTACs that target the tau protein^26,27^. bioPROTACs are potentially powerful tools for cell engineering because they can be easily designed to target different POIs via domain swapping and are genetically encodable, enabling their composition into genetic circuits.

Here, we introduce novel bioPROTACs optimized for T cell engineering. These bioPROTACs leverage a variety of different protein binding domains and utilize a minimal degron to degrade proteins. We then show that these optimized bioPROTACs retain their activity in other mammalian cell types. Using rational design strategies, we optimized a bioPROTAC capable of removing over 99% of a second-generation CAR from the plasma membrane. Furthermore, these bioPROTAC-inhibited CAR T cells exhibited substantially impaired cytotoxicity and proliferation in *in vitro* coculture models. Using one of our bioPROTACs, we built a circuit that degrades endogenous ZAP70 upon recognition of a specific surface antigen. ZAP70 is a cytoplasmic tyrosine kinase that is recruited to the plasma membrane and binds phosphorylated immunoreceptor tyrosine-based activation motifs (ITAMs) found in both the TCR complex and CARs. At the cellular level, ZAP70-deficiency results in inability to activate signaling events following TCR stimulation^30,31^. Furthermore, knocking out ZAP70 in CAR T cells results in failure to propagate downstream signals following CAR stimulation^32^. Taking advantage of the essential role of ZAP70, our bioPROTAC circuit degrades ZAP70 in response to intercellular interactions to weaken CAR T cell signaling. This work demonstrates the potential of bioPROTAC genetic circuits to modulate synthetic and endogenous signaling in therapeutic cell engineering.

## Results and Discussion

### Novel bioPROTACs potently degrade cytosolic proteins in T cells

Targeted protein degradation can be achieved by recruiting the ubiquitin proteasome system (‘UPS’) to a protein of interest via heterobifunctional molecules that combine a target recognition domain with a UPS recruiting domain. In this work, we optimized a TPD tool for cell engineering by minimizing the UPS recruitment domain in the form of a short peptide degron, enhanced activity through rational protein engineering, and demonstrated broad utility in human T cells (Fig 1A).

**Figure 1.**
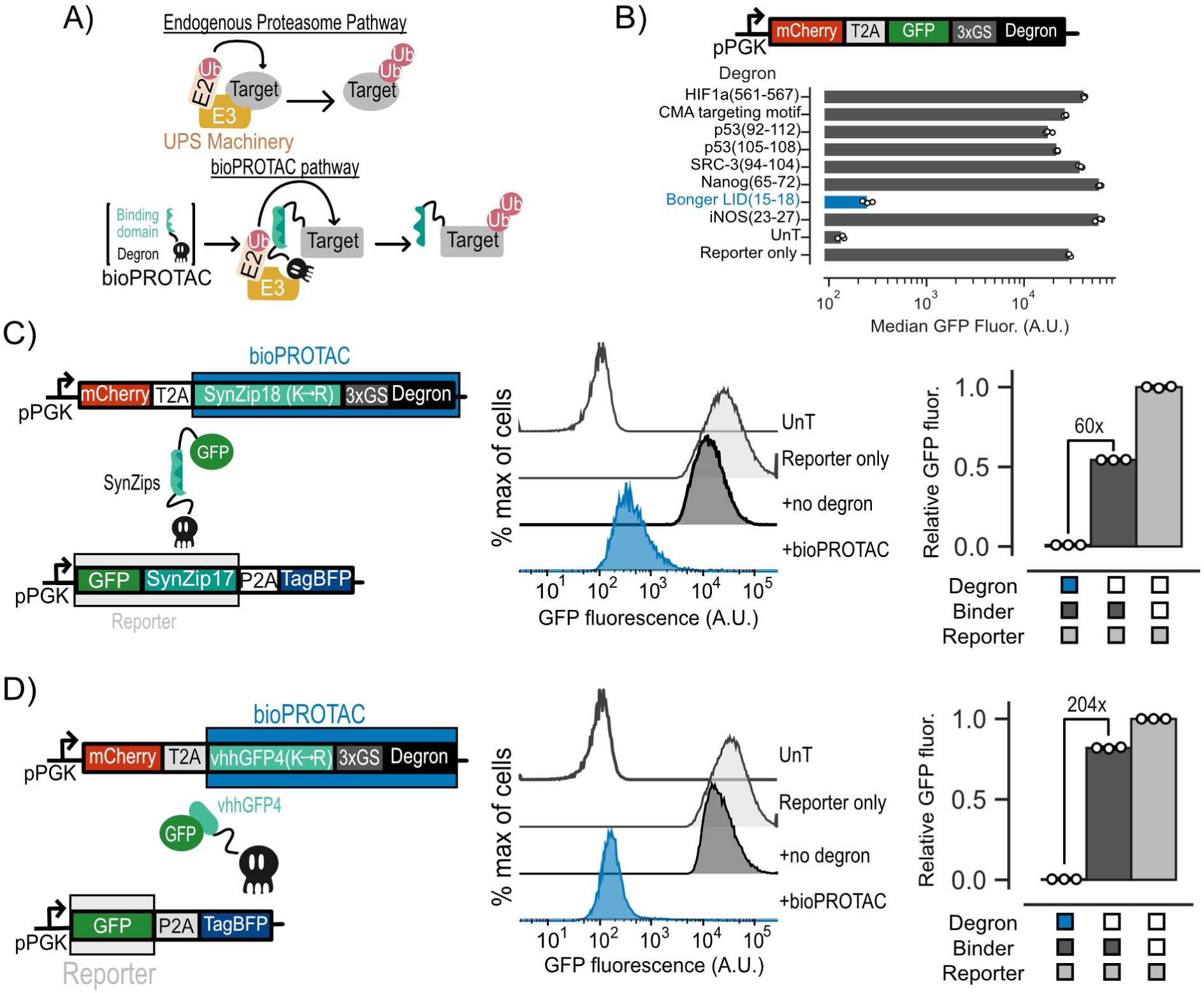
Novel bioPROTACs potently degrade cytosolic proteins in T cells. (A) Top: Cartoon diagramming abbreviated endogenous proteasomal degradation. Bottom: Cartoon depicting proposed bioPROTAC design and implementation. (B) Top: Cartoon depicting archetype of lentiviral payload used to engineer Jurkat T cells for screening of degrons for bioPROTAC construction. Bottom: Comparison of degron efficacies. Fluorescence was measured by flow cytometry. Dots represent biological replicates and error bars show SEM. (C and D) Left: Cartoons describing lentiviral payloads used to test bioPROTAC efficacy in Jurkat T cells. Middle: Flow cytometry histograms of the diagrammed lentiviral payloads in the left panel alongside controls. Histograms are representative of three independent experiments. Right: Quantification of flow cytometry histograms. Relative GFP fluorescence was calculated by normalizing measured GFP fluorescence by the fluorescence of the reporter only control. Each dot represents a technical replicate and error bars show SEM. Data is representative of three independent experiments.

The majority of existing bioPROTACs utilize truncated E3 ubiquitin ligases to recruit the UPS. This approach is not favorable for cell engineering because E3 ubiquitin ligases and their domains can be quite large, posing a potential challenge for delivery of large genetic payloads into the cell^28^. In this work, we explored bioPROTAC designs utilizing degrons to recruit the UPS because of their smaller genetic payload size. This is essential for our goal of constructing complex genetic circuits, as efficient genetic payload delivery in T cells is negatively correlated with payload size^33^. We first screened previously published degron sequences known to promote ubiquitination for degradation activity by fusing these degrons to the C-terminus of GFP^34–37^. We transduced Jurkat cells with individual lentiviral vectors encoding each candidate of this screen and measured GFP fluorescence by flow cytometry. We selected cells that were mCherry+ relative to an untransduced control for further analysis. From these data, we found that the minimal sequence responsible for protein degradation from the ligand induced degron (‘LID’) described by Bonger et al. induced over 116-fold decrease in GFP fluorescence relative to a reporter only control, the highest in our screen. By contrast, we did not observe a remarkable effect on GFP fluorescence from the other degrons in this screen (Fig 1B). The compact size of this sequence from the Bonger LID, comprising only four residues (RRRG), is ideal for applications in cell engineering. Future experiments in this work will use this RRRG sequence and simply refer to it as “degron”.

Next, we tested whether the degron could induce *trans* degradation of GFP as a model target protein. We hypothesized that bringing our degron and GFP in close proximity could promote ubiquitination and degradation of GFP through degron-based recruitment of endogenous E3 ubiquitin ligases. We designed several bioPROTACs exploring the use of the vhhGFP4 anti-GFP nanobody as well as a suite of synthetic leucine zippers (SynZips) as binders^38,39^. SynZips are a powerful tool for cell engineering due to their compact size, well-characterized affinity and large set of orthogonal binders. In this work, we explored the use of SynZip17 (cognate binder SynZip18) and SynZip1 (cognate binder SynZip2) for recruitment of the degron to our target protein. These binders were fused to the degron via a short, flexible 3x(GS) linker. As a “no degron” negative control, we substituted the degron in this design for a previously published four residue sequence ‘TRGN’ that demonstrates no degron activity^34^. To test our bioPROTACs, we co-transduced Jurkat T cells with one lentivirus encoding a GFP target protein and another lentivirus encoding a bioPROTAC. Recruitment of the SynZip bioPROTACs required tagging the GFP with a C-terminal fusion of the cognate SynZip. We then measured GFP fluorescence by flow cytometry. We selected cells that were mCherry+ and tagBFP+ relative to an untransduced control for further analysis. To assess degradation efficacy of each bioPROTAC variant, we calculated “Relative GFP fluorescence” which is the GFP fluorescence of each test condition normalized to the fluorescence of cells transduced with GFP alone. While bioPROTACs utilizing the SynZip binders exhibited strong degradation, the vhhGFP4 nanobody-based bioPROTAC showed only 1.45-fold reduction in GFP fluorescence (Fig S1).

We hypothesized that cis-ubiquitination could be limiting the activity of the vhhGFP4 nanobody bioPROTAC by inducing degradation of the bioPROTAC in addition to the POI. Previous work has shown that mutating lysine residues of a heterobifunctional protein degrader can improve its activity by minimizing cis-ubiquitination of the degrader and enhancing trans-ubiquitination of the POI^40^. We applied this logic to our initial designs by replacing lysine residues on each binding domain with arginine (‘K ->R mutation’). By performing the lysine substitution on the SynZip18 binder, we observed a 12-fold increase in activity over the wild-type binder, resulting in a 60-fold decrease in GFP fluorescence relative to the no degron control (Fig 1C). The same substitution on the vhhGFP4 nanobody binder resulted in a 100-fold increase in activity over the wild-type binder, resulting in a 204-fold decrease in GFP fluorescence relative to the no degron control (Fig 1D). The optimized bioPROTACs incorporating either the SynZip18 or vhhGFP4 nanobody with K->R mutation fused to the RRRG degron are the foundation for the subsequent experiments in this work.

### Our bioPROTACs are a versatile tool that function through the UPS and cullin ring ligases

To determine the wider applicability of these new bioPROTACs, we tested them in mouse embryonic stem cells (‘mESCs’), primary human CD4+ T cells (‘T cells’), 3T3 mouse fibroblasts, HEK293T cells and K562 leukemia cells by co-transducing a GFP reporter and either a vhhGFP4 nanobody or SynZip18 based bioPROTAC via lentivirus. We selected cells that were mCherry+ and tagBFP+ relative to an untransduced control for further analysis. We found that the SynZip18 bioPROTAC reduced GFP fluorescence by at least 33-fold relative to reporter only controls in all tested cell types, demonstrating the versatility of our bioPROTAC for cell engineering (Fig 2A). The vhhGFP4 nanobody bioPROTAC displayed similar activity in these cell types (Fig S2A).

**Figure 2.**
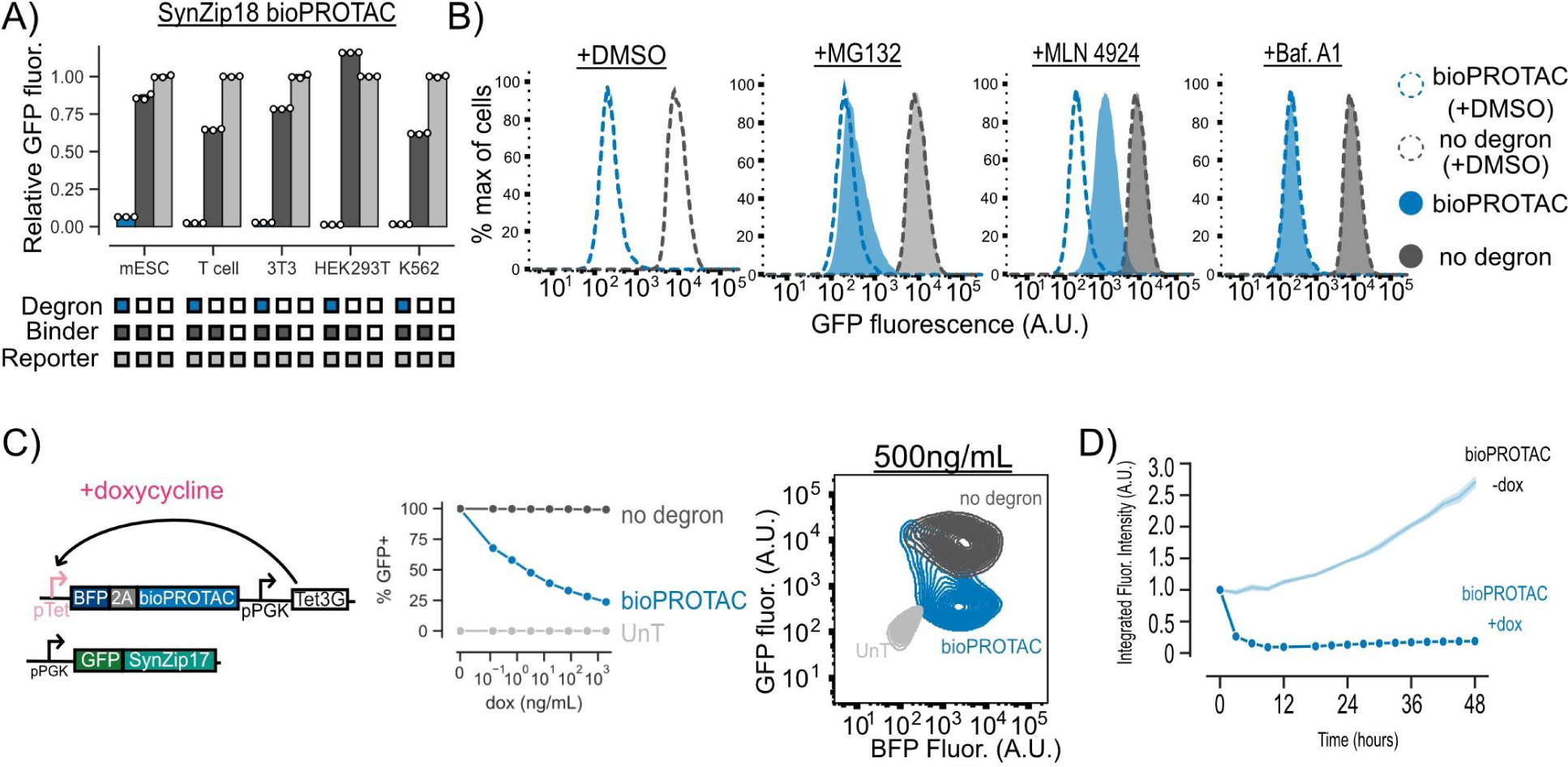
bioPROTACs are a versatile tool for protein degradation capable of dose-dependent and rapid degradation of proteins. (A) bioPROTACs are capable of potent degradation across a variety of mammalian cell types. GFP fluorescence was measured by flow cytometry and normalized to a GFP reporter alone for each cell type. Each dot represents a technical replicate and error bars show SEM. (B) bioPROTAC degradation of cytosolic proteins relies on the proteasome via cullin ring ligases. SynZip18 bioPROTAC and GFP reporter expressing Jurkat T cells or control lines were treated with either MG132, Bafilomycin A1, MLN4924 or a DMSO control. GFP fluorescence was then measured by flow cytometry. Histograms are representative of three biological replicates. (C) bioPROTACs titration results in dose-dependent degradation of cytosolic proteins. Left: Cartoon depicting lentiviral payloads encoding a GFP reporter protein and a doxycycline inducible bioPROTAC and the Tet3G protein. Middle: After isolation by FACS, cells were treated with a 2-fold titration series of dox or a media only control for 48 hours. bioPROTAC efficacy was assessed by flow cytometry. Each dot represents the mean of three biological replicates. Error shows SEM. Right: Representative contour plot of bioPROTAC expression level (BFP) and GFP expression level at 500 ng/mL of doxycycline. (D) bioPROTACs induced substantial GFP loss after 4 hours of doxycycline treatment. The doxycycline inducible Jurkat T cell line described above was treated with either 2000 ng/mL doxycycline or a media only control. GFP fluorescence was measured by live cell imaging and analyzed using the Incucyte software. Each trace is normalized to integrated fluorescent intensity values at the initial measurement time point. Each dot is the mean of three biological replicates. Error shows SEM.

To explore the mechanism of GFP degradation via bioPROTACs, we used a small molecule panel to interrogate proteasomal and lysosomal pathways. Jurkat T cells expressing GFP and the SynZip18 bioPROTAC were treated with either the proteasomal inhibitor MG132, the vacuolar ATPase inhibitor Bafilomycin A1, the pan cullin ring E3 ligase inhibitor MLN4924, or a DMSO control for 5 hours. After treatment, cells were washed and GFP fluorescence was measured by flow cytometry. We selected cells that were mCherry+ and tagBFP+ relative to an untransduced control for further analysis. We observed minor rescue of GFP fluorescence in cells treated with MG132 and much higher levels of rescue in cells treated with MLN4924 relative to a DMSO vehicle control. These data suggest that the bioPROTAC-dependent loss of GFP fluorescence is due to protein degradation through the UPS mediated by cullin ring ligases. By contrast, GFP fluorescence was minimally affected in cells treated with Bafilomycin A1, suggesting minimal contribution of the lysosome to bioPROTAC activity. In order to further investigate the mechanism of bioPROTAC-dependent GFP loss, we co-expressed a series of dominant negative cullin constructs to inhibit endogenous cullin function^41^. The dominant negative cullins are C-terminally truncated mutants of cullins and retain the ability to bind their targets but are unable to promote ubiquitin conjugation, thereby blunting degradation through specific cullin pathways. We observed a rescue of GFP fluorescence when the bioPROTAC was co-expressed with a dominant negative cullin from the cullin 4A and 4B families, suggesting the RRRG degron is recognized by a a cullin ring ligase belonging to these families of cullins in T cells (Fig S2D).

### Kinetics of bioPROTAC-mediated degradation

We then explored the relationship between bioPROTAC expression level and bioPROTAC-mediated degradation. Jurkat T cells were engineered to express a constitutive GFP reporter protein and doxycycline-inducible expression of the SynZip18 bioPROTAC. We titrated SynZip18 bioPROTAC expression with a serial dilution of doxycycline and measured GFP fluorescence via flow cytometry after 48 hours of treatment. We observed a decreasing number of GFP expressing cells with higher concentrations of doxycycline, indicating dose-dependent SynZip18 bioPROTAC activity. To better understand the relationship between SynZip18 bioPROTAC concentration and activity, we plotted GFP fluorescence versus BFP fluorescence (BFP being a proxy for SynZip18 bioPROTAC expression) for cells treated with 500 ng/mL doxycycline. At intermediate BFP expression levels, the GFP signal was bimodal, with a small number of cells that retained high GFP expression and a majority of cells that had lost GFP expression. At high BFP expression levels, GFP fluorescence was completely lost, indicating SynZip18 bioPROTAC activity is thresholded on expression level (Fig 2C, Fig S2B). The vhhGFP4 nanobody bioPROTAC exhibited similar dose-dependent degradation activity under similar experimental conditions (Fig S2C).

In order to explore the kinetics of SynZip18 bioPROTAC-mediated degradation, we next treated cells with a saturating dose of 2000 ng/mL doxycycline and measured GFP intensity over 48 hours using fluorescence microscopy and live cell imaging. We observed loss of GFP fluorescence within 3 hours, and complete degradation of GFP within 12 hours while the untreated control demonstrated increasing GFP fluorescence over the course of the experiment (Fig 2D, Fig S2B). The kinetics of SynZip18 bioPROTAC degradation are comparable to that of previously published bioPROTACs, but slower than other published protein-based degraders that can pre-complex with their POI, such mAID-nanobody, prior to activation^18,40^.

### Plasma membrane localization of bioPROTACs enhances internalization of CAR

We next set out to test whether our bioPROTACs could induce internalization of CAR from the plasma membrane. Jurkat T cells were lentivirally transduced with a single plasmid encoding a GFP transduction marker, a SynZip18 bioPROTAC, and a CAR separated by 2A elements (Fig 3A). To recruit the bioPROTAC, we fused SynZip17 to the C-terminus of the CAR. For simplicity, we will still refer to this modified fusion molecule as a CAR. We tested bioPROTAC-induced internalization of two CAR variants, an anti-CD19 4-1BBz CAR and an anti-HER2 4-1BBz CAR, from the plasma membrane. We used flow cytometry to measure surface expression of the CAR molecules after immunofluorescence staining for a myc epitope tag fused to the extracellular domain of the CAR. We selected cells that were GFP+ relative to an untransduced control for further analysis. To quantify SynZip18 bioPROTAC activity, we calculated the percentage of CAR remaining at the cell surface by subtracting background stain fluorescence, and normalizing to the CAR only control for each tested condition. The bioPROTAC reduced anti-CD19 CAR fluorescence by 80%, but exhibited minimal effect on the anti-HER2 CAR (Fig 3B, Fig 3C). Despite the reduction in anti-CD19 CAR fluorescence, substantial anti-CD19 CAR expression was still detectable at the cell surface. Previous work has demonstrated that low levels of CAR expression in CAR T cells can still result in complete clearance of tumor cells *in vitro*^42^. We therefore sought to maximize SynZip18 bioPROTAC-mediated internalization of CAR at the plasma membrane and minimize any potential signaling.

**Figure 3.**
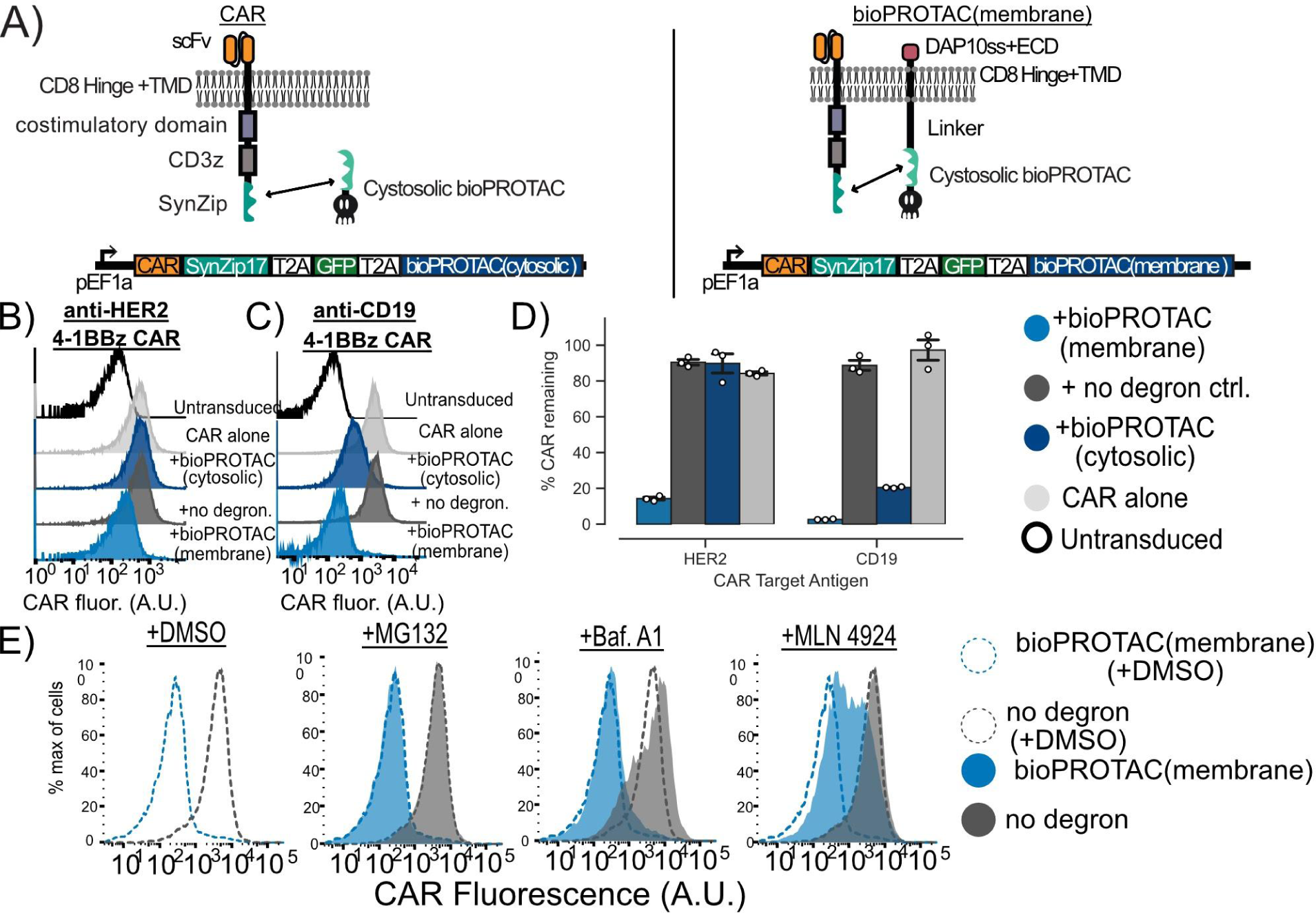
Plasma membrane localization of bioPROTACs enhances internalization of CAR. (A) Cartoons diagramming lentiviral payloads for targeting CARs with bioPROTACs. (B) Membrane tethered bioPROTAC (‘bioPROTAC(membrane)’) is capable of internalization of a second-generation anti-HER2 4-1BBz CAR. Flow cytometry histograms using the plasmids demonstrating knockdown of anti-HER2 CAR in Jurkat T cells. Histograms are representative of three technical replicates. (C) SynZip18 bioPROTAC(membrane) is capable of potent internalization of anti-CD19 4-1BBz CAR with some effect from the bioPROTAC(cytosolic). Flow cytometry histograms representing Jurkat T cells expressing the plasmids diagrammed in Figure 3A. Histograms are representative of three technical replicates. (D) Quantification of flow cytometry data. Flow cytometry data is normalized to CAR alone controls and quantified. Each dot here represents technical replicates. (E) Cullin ring ligases are implicated in CAR internalization by bioPROTAC(membrane), but not the proteasome. The anti-CD19 CAR Jurkat T cells lines described above were treated with the same panel of drugs described in previous experiments. Cells were then washed and CAR fluorescence was measured by antibody staining and flow cytometry. Histograms are representative of three biological replicates.

We hypothesized that increasing the concentration of the SynZip18 bioPROTAC at the plasma membrane could mediate more efficient internalization of CAR from the plasma membrane by enhancing colocalization. We fused either a previously described chimeric DAP10 signal sequence-DAP10 extracellular domain-CD8a transmembrane domain, or a lyn membrane targeting tag to the SynZip18 bioPROTAC via a rigid 15 amino acid linker to create a new molecule that we refer to as the membrane tethered bioPROTAC or ‘bioPROTAC(membrane)’^43,44^. We accordingly renamed the original non-tethered design the “bioPROTAC(cytosolic)”. We compared the activity of the SynZip18 bioPROTAC(cytosolic), the SynZip18 bioPROTAC(membrane) and a no degron control for the SynZip18 bioPROTAC(membrane) where the degron is swapped for the same control peptide sequence used previously. We tested our new SynZip18 bioPROTAC(membrane) against the CARs as before and compared its activity to the SynZip18 bioPROTAC(cytosolic). When targeting an anti-HER2 CAR, the SynZip18 bioPROTAC(membrane) removed 86% of detectable CAR relative to the CAR alone control, a substantial improvement over the SynZip18 bioPROTAC(cytosolic) (Fig 3B, Fig 3D). Similarly, the SynZip18 bioPROTAC(membrane) augmented internalization of the anti-CD19 CAR, resulting in over 99% loss of the anti-CD19 CAR fluorescence relative to the CAR alone control (Fig 3C, Fig 3D). The SynZip18 bioPROTAC(membrane) was also effective against anti-CD19-CD28z and anti-HER2-CD28z CARs, but showed weaker levels of activity than against anti-CD19 4-1BBz and anti-HER2 4-1BBz CARs (Fig S3A). We observed similar loss in anti-CD19 4-1BBz CAR fluorescence from the lyn tag variant of the SynZip18 bioPROTAC(membrane), suggesting that various strategies for membrane targeting can enhance bioPROTAC activity against CARs (Fig S3B). We decided to continue future experiments with the DAP10 localized bioPROTAC(membrane), because bioPROTAC(membrane) expression can be detected through antibody staining for a V5 peptide tag fused to the DAP10 extracellular domain (Fig S3C).

In order to investigate the mechanism of bioPROTAC(membrane)-mediated CAR internalization, we treated Jurkat cells expressing the anti-CD19 4-1BBz CAR and the SynZip18 bioPROTAC(membrane) with the previously described proteasomal and lysosomal inhibitors and measured CAR expression by flow cytometry. CAR expression was rescued by MLN4924 and unaffected by Bafilomycin A1. In contrast to the bioPROTAC(cytosolic) against GFP, MG132 had minimal effect on SynZip18 bioPROTAC(membrane)-mediated CAR internalization from the plasma membrane (Fig 3E). This result suggests that loss of detectable CAR expression from the plasma membrane may be mediated through cullin ring ligases, but not the proteasome. From these data, we were able to conclude that the SynZip18 bioPROTAC(membrane) internalizes CARs from the plasma membrane in a model of human T cells.

### Membrane tethered bioPROTAC abrogates CAR T cell signaling in primary human T cells

Next, we sought to test whether the levels of bioPROTAC-mediated CAR internalization observed in Jurkat cells would translate to functional consequences in primary human CAR T cells. We transduced primary CD8+ human T cells with a single lentivirus encoding the SynZip18 bioPROTAC(membrane), anti-CD19 CAR, and GFP separated by 2A elements. To control for the effect of SynZip18 bioPROTAC(membrane) binding on CAR function, we also engineered T cells with the no degron SynZip18 bioPROTAC(membrane) negative control.

We challenged these anti-CD19 CAR T cells with K562 leukemia cells expressing either CD19 or no antigen at a 1:1 effector to target (‘E:T’) ratio. After 72 hours of co-culture, we measured CAR T cell cytotoxicity and CAR expression via flow cytometry. In concordance with data collected in Jurkat cells, the SynZip18 bioPROTAC(membrane) removed over 99% of detectable surface expressed CAR in primary T cells (Fig 4A). This level of SynZip18 bioPROTAC(membrane)-mediated CAR internalization translated to a mere 6% lysis of the antigen expressing cell population by CAR T cells in co-culture. In comparison, the no degron control CAR T cells lysed 66% of the antigen expression cell population in co-culture (Fig 4B). In the same assay, we explored the effect of bioPROTAC(membrane)-mediated CAR internalization on the activation of CD25, a canonical marker for T cell activation. CD25 is a key regulator for activation-related T cell proliferation as part of the high affinity IL-2 receptor and has shown to be upregulated in response to CAR signaling^45–48^. SynZip18 bioPROTAC(membrane)-mediated internalization of CAR heavily downregulated expression of CD25 in engineered T cells resulting in 71% fewer engineered T cells expressing CD25 relative to the no degron control and a mere 10% increase in CD25+ cells relative to untransduced primary human T cells (Fig 4C). These results suggest that the SynZip18 bioPROTAC(membrane) is able to achieve a functionally relevant level of internalization of CAR resulting in inhibition of multiple facets of CAR signaling.

**Figure 4.**
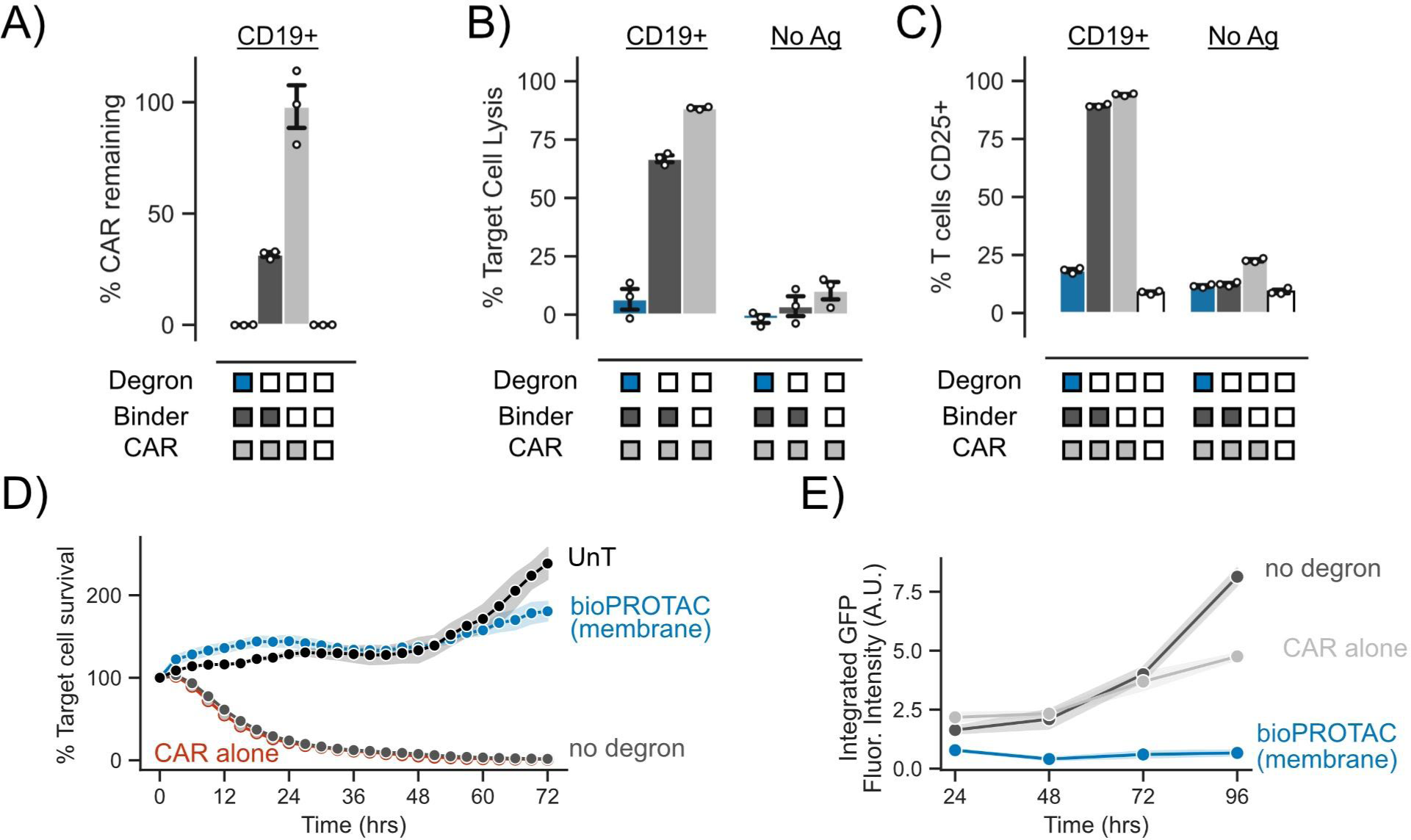
Membrane tethered bioPROTAC abrogates CAR T cell signaling in primary human T cells. (A) bioPROTAC(membrane) co-expression reduced CAR expression in primary T cells. ‘% CAR remaining’ is background subtracted measured CAR fluorescence normalized to the CAR only control. Dots represent technical replicates and error shows SEM. (B) bioPROTAC(membrane) expression completely inhibits CAR cytotoxicity. Using flow cytometry, we measured CAR T cell cytotoxicity in a 1:1 E:T ratio using CD19 expressing or no antigen K562s. Dots represent technical replicates and error shows SEM. (C) bioPROTAC(membrane) prevented upregulation of T cell activation marker CD25. CD25 levels were measured by immunostaining followed by flow cytometry. Dots represent technical replicates and error shows SEM. (D) bioPROTAC(membrane) ablated CAR cytotoxicity even at high E:T ratios sampling during a 72 hour time course. Engineered T cell lines described above and challenged them with Nalm6 cells at an E:T ratio of 3:1 and assayed for target cell clearance by live cell microscopy. Each dot represents the mean of three technical replicates and error shows SEM. (E) bioPROTAC(membrane) stifled CAR T cell proliferation and survival. In this same assay as described in Figure 4D, we observed T cell population numbers by GFP fluorescence over 72 hours by Incucyte image analysis. Each dot represents the mean of three technical replicates and error shows SEM.

Following these results, we explored how the SynZip18 bioPROTAC(membrane) affects CAR induced cytotoxicity and CAR T cell population levels over time by utilizing a previously published model of mCherry expressing Nalm6 acute lymphoblastic leukemia cells^49^. We increased the E:T ratio for these experiments to 3:1 to understand the extent of CAR signaling inhibition by the SynZip18 bioPROTAC(membrane) when T cells are in an overwhelming excess to the tumor cells. We measured the growth of Nalm6 populations in these co-culture conditions by live cell fluorescence imaging of mCherry every 3 hours for 72 hours. Gratifyingly, SynZip18 bioPROTAC(membrane) co-expressing CAR T cells exhibited no observable cytotoxicity against Nalm6 cells and behaved similarly to untransduced T cells. By contrast, CAR T cells co-expressing the no degron control lysed 95% of the Nalm6 population within 24 hours relative to the initial time point (Fig 4D, Fig S4A).

We simultaneously tracked CAR T cell populations by quantifying GFP fluorescence levels using live cell fluorescence imaging for 72 hours. CAR T cell cells co-expressing the SynZip18 bioPROTAC(membrane) demonstrated no observable proliferation relative to the initial time point, whereas the no degron control expressing CAR T cells showed a 2.4-fold increase in CAR T cell numbers over the course of the experiment (Fig 4E). Together, these data demonstrate that the SynZip18 bioPROTAC(membrane) functionally inhibits multiple facets of CAR signaling in primary T cells. We observed similar SynZip18 bioPROTAC(membrane)-dependent suppression of the anti-HER2 CAR, suggesting that SynZip18 bioPROTAC(membrane)-mediated knockdown may be a generally effective strategy for inhibiting CAR signaling (Fig S4B-D). We also observed that the weaker internalization of the CAR by the SynZip18 bioPROTAC(cytosolic) resulted in lysis of over 25% of the target cell population (Fig S4E).

### bioPROTACs can be composed into circuits for cell autonomous modulation of CAR T cell signaling

A key characteristic of bioPROTACs is their ability to be genetically encoded. This trait enables the construction of genetic circuits that utilize bioPROTACs to modulate cellular signaling in response to environmental cues, independently of exogenous input such as small molecules. As a proof of concept, we designed a genetic circuit to disrupt CAR T cell signaling upon recognition of a user-defined antigen. To achieve signaling disruption, we chose to target an essential component of the TCR signaling pathway, the tyrosine kinase ZAP70, for degradation. To enable simultaneous recruitment of the vhhGFP4 nanobody bioPROTAC to ZAP70 and target protein tracking, we adapted a previously published method utilizing CRISPR/Cas9 to tag the N-terminus of endogenous ZAP70 with GFP in primary human CD4+ T cells^50^. After genome engineering, we isolated the GFP+ population via FACS (Fig 5A). We further engineered these T cells via lentiviral transduction of an anti-CD19 4-1BBz CAR and an anti-HER2 SNIPR that activates expression of the vhhGFP4 nanobody bioPROTAC upon encountering a target antigen. Based on this circuit design, SNIPR binding to the target antigen HER2 should trigger expression of the bioPROTAC and degrade GFP-ZAP70 (Fig 5B). To test the functionality of our circuit, we cocultured our engineered T cells with K562 target cells expressing either CD19, or CD19 and HER2. After 48 hours of coculture, we performed flow cytometry to measure GFP expression. Gratifyingly, we observed that 47% fewer engineered T cells expressed GFP when expressing the intact bioPROTAC circuit in the presence of HER2 expressing target cells. By contrast, GFP expression in T cells expressing a no degron negative control bioPROTAC was unaffected by the presence of HER2 (Fig 5C).

**Figure 5.**
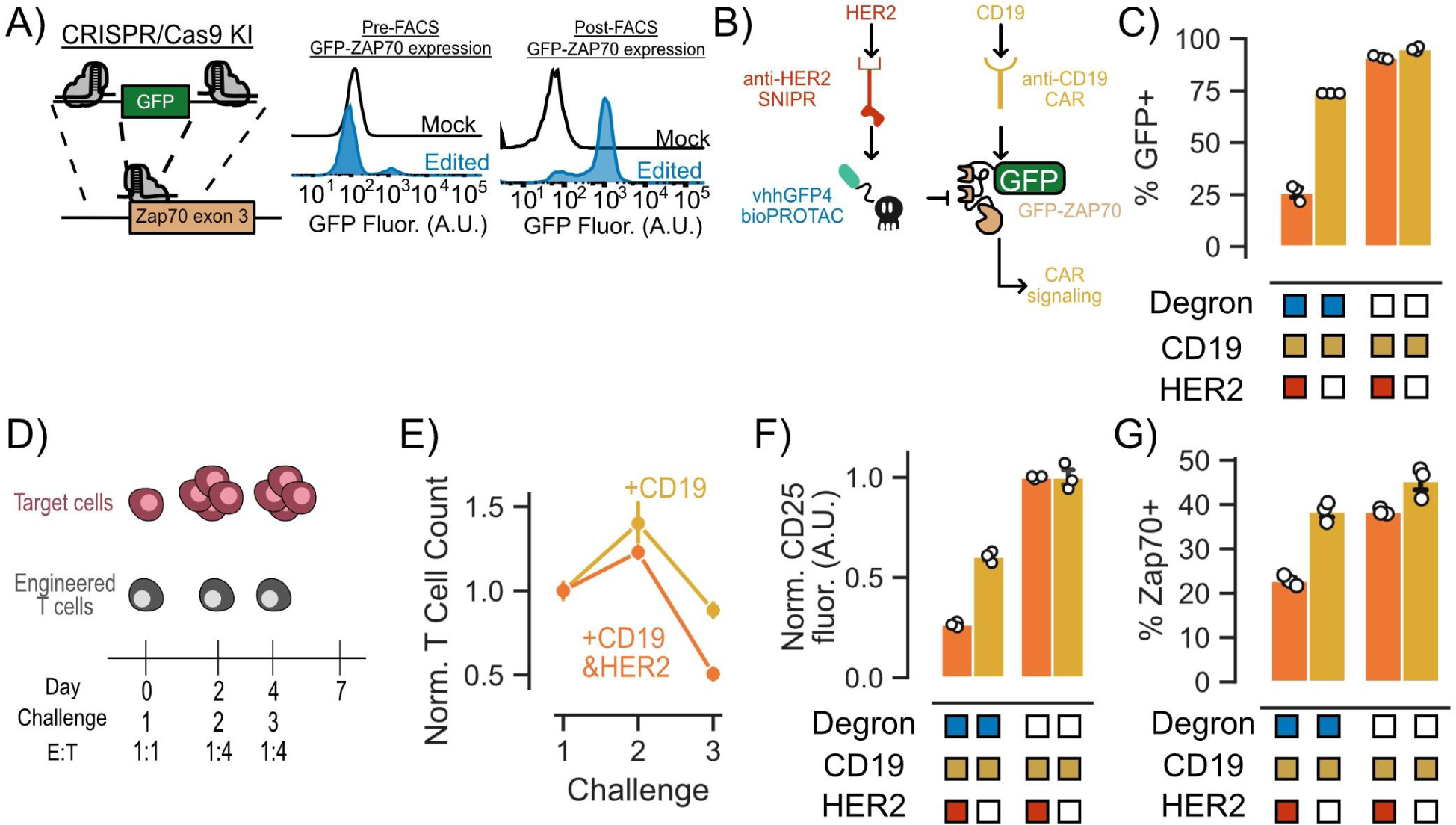
bioPROTACs can be composed into circuits for cell autonomous modulation of CAR T cell signaling. (A) CRISPR/Cas9 knock-in technology can be used to guide bioPROTACs to endogenous proteins of interest. (B) Genetic circuit uses SNIPR antigen detection to trigger bioPROTAC expression resulting in GFP-ZAP70 degradation. (C) SNIPR-induced bioPROTACs induced degradation of GFP-ZAP70 as measured by GFP fluorescence. Engineered T cells were challenged with K562 target cells expressing CD19 or CD19 and HER2 at an E:T ratio of 1:1 for 48 hours. GFP fluorescence was measured by flow cytometry. (D) Multiple challenge assay used to assess circuit functionality. (E) T cell proliferation and survival is affected by bioPROTAC mediated knockdown of GFP-ZAP70. T cell numbers were quantified using flow cytometry and normalized to the no degron control for each antigen condition. T cell counts were normalized to the no degron control and then again normalized to the first challenge. Dots are the mean of three technical replicates and error bars represent SEM. (F) bioPROTAC-mediated knockdown of GFP-ZAP70 resulted in functional consequences in CAR T cell signaling. Each dot on the line represents the mean of three technical replicates. The error shown represents the SEM. CD25 fluorescence values were normalized to those of cells expressing the no degron control circuit. (G) SNIPR-induced bioPROTACs are capable of inducing GFP-ZAP70 degradation as measured by total ZAP70 levels. ZAP70 fluorescence was measured by intracellular staining followed by flow cytometry. ZAP70+ cells were determined by *in silico* gating. Each dot represents technical replicates and error bars show SEM. ZAP70 fluorescence values were normalized to those of cells expressing the no degron control circuit.

To explore the effect of the bioPROTAC genetic circuit on engineered CAR T cells over time, we designed a repeated challenge assay where we subjected T cells to a 1:1 E:T ratio coculture for 48 hours, followed by a second and third round of coculture at a 1:4 E:T ratio at days 2 and 4 (Fig 5D). Considering the time delay between antigen-dependent activation of bioPROTAC expression, degradation of ZAP70, and signaling outcomes, we hypothesized that the phenotypic effect of our circuit may not be evident within the duration of a single tumor cell challenge. Indeed, 48 hours after the first challenge, we observed a decrease in GFP-ZAP70 levels, but detected no change in CD25 expression or T cell proliferation for engineered T cells cultured with CD19 and HER2 versus CD19 alone target cells (Fig 5E, Fig S6). After a second round of co-culture, we observed a 16% reduction in T cell proliferation and survival for T cells co-cultured with CD19 and HER2 target cells compared to T cells co-cultured with CD19 alone target cells. This difference became more pronounced after a third tumor cell challenge, with bioPROTAC circuit-expressing CAR T cells showing a 45% reduction in proliferation and survival in the presence of HER2 relative to circuit-expressing CAR T cells cultured in the absence of HER2 (Fig 5E). We also observed that the bioPROTAC circuit reduced CD25 expression by 50% in the presence of the HER2 when compared to CD25 expression of circuit-expressing cells in the absence of HER2 (Fig 5F). To further validate the role of the bioPROTAC circuit in CAR T cell modulation, we explored how activation of the circuit affected levels of total ZAP70 using intracellular staining and flow cytometry. In concordance with the phenotypic data, we observed 16% fewer ZAP70+ cells in bioPROTAC circuit expressing cells in the presence of HER2 than in the absence of HER2 (Fig 5G). With these data, we demonstrated that bioPROTACs can be composed into a genetic circuit that degrades a critical signaling protein in an antigen-dependent fashion to modify CAR T cell phenotypes.

## Conclusions

In order to create an ideal bioPROTAC for T cell engineering, we drew inspiration from the endogenous UPS and previously published bioPROTACs. We had two major design requirements: compactness and POI targeting modularity. Here, we introduced a bioPROTAC as small as 55 amino acids, making it ideal for delivery for cell engineering and for use in genetic circuits. We demonstrated the ability of the Bonger LID minimal degron sequence, RRRG, to be composed into several bioPROTACs with structurally distinct protein binding domains. This degron has also previously been utilized in a bioPROTAC targeting α-Synuclein, further supporting the modularity of this UPS-recruitment approach. Exploring other protein binding domains as binders for the degron-based bioPROTAC may hold promise for future engineering, therapeutic, and cell biology applications. Furthermore, advancements in *de novo* protein design and antibody engineering are driving forward the possibility of designing binders for any protein of interest^51,52^.

Recent work has demonstrated that bioPROTACs (referred to as SySRs) can enhance CAR T cell potency by degrading SMAD2 and SMAD3, thus blocking SMAD-dependent TGFβ signaling^53^. SySR’s use of an endogenous protein as a binder highlights the potential to leverage other endogenous PPIs in bioPROTAC designs. The protein engineering strategies detailed in our work as well as the potent RRRG degron have the potential to enhance the activity of the SySR bioPROTAC. A future cell therapy could coordinate the bioPROTACs used in our work with the SySR bioPROTAC to balance the ablation and augmentation of CAR T cell-related signaling for more sophisticated cell autonomous tumor control.

We also found that activity of bioPROTACs can be tuned through mutation of lysine residues in the binding domain. Interestingly, we found that mutagenesis improved SynZip18 and vhhGFP4 nanobody bioPROTAC degradation, but had minimal effect on the SynZip2 based bioPROTACs. We hypothesize that this could be attributed to availability of surface lysines of each binder. Further investigation into how these structurally similar proteins are differentially affected by lysine substitution could be an interesting avenue for optimizing future bioPROTAC designs and give insights into the mechanisms behind cis-ubiquitination.

Another useful property of our bioPROTACs is their robust activity across many mammalian cell types. We show that our bioPROTACs function in commonly used cell lines, such as HEK293T cells, which can be useful for prototyping potential circuit architectures and guiding foundational cell engineering principles. We also show that bioPROTACs are an effective degrader in mESCs, which are used as a classic model in developmental biology and synthetic development biology. While engineered T cells are immensely successful in the clinic, other cell types, including human stem cells, natural killer (NK) cells and macrophages, have shown great promise. Future exploration of bioPROTACs in these cell types could help unlock new therapeutic applications^54,55^. Upon investigation into the degradation mechanism of our bioPROTACs, we identified cullin ring ligases as playing a key role in the degradation of both cytosolic and membrane proteins in Jurkat cells. Specifically, we identified the CUL4A and 4B families as mediators of bioPROTAC-based degradation, whereas perturbing the CUL2 family did not have an observable effect. This conflicts with previously described “rules” of C-terminal degron degradation of the mammalian proteome^41^. We hypothesize that this discordance may hint at cell-type specific preferences for cullin families in UPS protein recognition. Further experiments to genetically knockdown or knockout specific E3 ligases would help elucidate the specific mechanism of degradation of our bioPROTACs.

We showed that bioPROTAC-based degradation of cytosolic protein targets utilizes the proteasome and relies on cullin ring ligases. We observed higher levels of rescue from treatment of Jurkat cells with MLN4924 than with MG132. We hypothesize this phenomenon may be attributed to MG132-induced apoptosis of Jurkat T cells due to ER stress^56^. While this may not impact the conclusions drawn from the data, future exploration into bioPROTAC use and development in Jurkat cells could optimize dosage and timing of MG132 treatment to yield better efficacy. In contrast to bioPROTAC-mediated degradation of cytosolic proteins, inhibition of the proteasome appeared to have no effect on a transmembrane protein target. This result replicated previous findings showing the inability of MG132 to prevent CAR internalization following antigen stimulation^57^. Interestingly, our data did not implicate the lysosome in the bioPROTAC-based internalization of membrane proteins, which contrasts with studies of other tools for targeting membrane proteins such as LYTACs and AbTACs^20,58^. However, we did observe rescue of CAR expression following cullin ring ligase inhibition. From these data we can conclude that ubiquitination is necessary for bioPROTAC(membrane) function, but the exact mechanism of bioPROTAC(membrane)-mediated CAR internalization is unclear. Further investigation into the mechanism of bioPROTAC-mediated internalization of membrane proteins may elucidate some unexpected degradation pathways.

Characterization of degradation kinetics showed that our bioPROTAC performs similarly to other previously published bioPROTACs. These measurements may assist in future efforts to build computational models of bioPROTAC circuits to guide circuit design. The use of post-translational methods to activate the bioPROTAC could vastly improve the kinetics of degradation. As a proof of concept, we replaced the transcription factor domain of an anti-HER2 SNIPR with our SynZip18 nanobody bioPROTAC (Fig S6A). This experiment highlights the potential of bioPROTACs to be coupled with existing post-translational control technologies to dictate bioPROTAC expression, activity, and localization^59–61^.

We improved bioPROTAC activity against membrane-bound target proteins through N-terminal fusion of a membrane tethering domain to the bioPROTAC. We found that this modification augmented bioPROTAC-mediated internalization of the antiHER2 4-1BBz CAR and resulted in complete removal of antiCD19 4-1BBz CAR from the plasma membrane. Unexpectedly, we observed that the no degron control for the bioPROTAC(membrane) decreased CAR fluorescence levels at the cell membrane in both Jurkat and primary T cells. While we observed a much greater level of CAR internalization with bioPROTAC(membrane), further exploration into whether this loss in fluorescence is due to steric occlusion of the epitope tag used to detect the CAR or if the binding of the SynZip pair is contributing to CAR internalization could help guide future bioPROTAC designs and applications.

To demonstrate the cell engineering potential of our bioPROTACs, we devised a circuit that uses environmental cues to activate bioPROTAC-mediated degradation of a key signaling protein in T cells. While our circuit was able to knockdown ZAP70 and reduce T cell proliferation, this strategy was insufficient to ablate target cell lysis (data not shown). This may be due to CAR signaling through non-ZAP70 mediated pathways. Direct degradation of the CAR using bioPROTACs could potentially circumvent this issue. However, our attempts to degrade the CAR via a SNIPR-induced bioPROTAC were also unsuccessful (data not shown). We hypothesize that this inability to regulate the CAR is due to an insufficient level of bioPROTAC expression from SNIPR activation. Since the CAR is expressed from a strong constitutive promoter, the CAR is likely present at a much higher concentration in the cell compared to endogenous signaling proteins such as ZAP70 and thus requires a higher concentration of bioPROTAC to be fully degraded.

Further optimization of circuits utilizing our bioPROTAC will be necessary before finding clinical applications. Given the time delay between SNIPR activation and bioPROTAC-induced downregulation of CAR-associated signaling, bioPROTACs may not be a suitable tool for building NOT logic gates for selective target cell recognition and killing. However, existing CAR T cell therapies suffer from CRS associated toxicity and T cell exhaustion induced hypofunctionality, both of which are phenomena that occur on the timescale of days or even weeks^62–64^. A circuit or promoter designed to sense high levels of IL-6, which is associated with the onset of cytokine release syndrome, could be used to activate expression of a bioPROTAC to downregulate T cell signaling and preempt the onset of CRS^65^. Similarly, a circuit or promoter designed to sense exhaustion-associated epigenetic changes could be used to degrade the CAR and enforce a period of rest, which has been shown to enhance CAR T cell functionality in *in vivo* models^6,66^. The compact nature of our bioPROTACs enables the potential delivery of these circuits in the form of a single lentivirus, or will greatly facilitate genomic integration.

More generally, genetic circuits that endow cell autonomous control over therapeutic functions could greatly improve the safety, specificity and efficacy of engineered cell therapies. Targeted protein degradation is a highly effective method for modulating signaling and complements current approaches in genetic circuit design. We believe that the bioPROTACs developed in this work will be useful for many future cell engineering endeavors.

## Materials and Methods

### Gene synthesis and cloning

All plasmids were cloned using the Mammalian Toolkit (MTK), a Golden-Gate based cloning system^67^. All parts plasmids from the MTK used in this work were domesticated from DNA sequences generated by oligonucleotide annealing, gBlocks or PCR. SNIPR plasmids were domesticated for use in the MTK by PCR using plasmids gifted by Dr. Kole Roybal as a template. Plasmids synthesized in this manner were propagated in Stbl3 *E. coli* from QB3 Macrolab. All part plasmids were verified by sequencing and all subsequent plasmids were verified by test restriction digest and/or sequencing.

### Jurkat T cell culture conditions

Jurkat T cells were cultured in RPMI-1640 (Gibco #11875-093) supplemented with 10% fetal bovine serum (FBS) and 1% anti-anti (Gibco #15240-096). Cells were maintained in T75 flasks and split 1:10 every 3 days.

### K562 target cell generation and cell culture conditions

K562 target cells expressing CD19, HER2 or CD19&HER2 were generated by lentiviral transduction using plasmids gifted by Dr. Wendell Lim and his lab. K562s were cultured in IMDM supplemented with 10% FBS and 1% gentamicin. K562 cells were maintained in T25 flasks and split 1:10 every 3 days.

### Source of primary human T cells

Blood was obtained from Blood Centers of the Pacific (San Francisco, CA) as approved by the University Institutional Review Board. Primary CD4+ and CD8+ T cells were isolated from anonymous donor blood after apheresis as described below.

### Primary human T cell isolation

Primary CD4+ and CD8+ T cells were isolated from anonymous donor blood after apheresis by negative selection. T cells were cryo-preserved in CellBanker cell freezing media.

### Cell culture for Lenti-X 293T cells

Lenti-X 293T packaging cells (Clontech #11131D) were cultured in medium consisting of Dulbecco’s Modified Eagle Medium (DMEM) (Gibco #10569-010) and 10% fetal bovine serum (FBS) (University of California, San Francisco Cell Culture Facility). Lenti-X 293T cells were cultured in T150 or T225 flasks (Corning #430825 and #431082) and passaged upon reaching 80% confluency. To passage, cells were treated with TrypLE express (Gibco #12605010) at 37 C for 5 minutes. Then, 10 mL of media was used to quench the reaction and cells were collected into a 50 mL conical tube and pelleted by centrifugation (400xg for 4 minutes). Cells were cultured until passage 30 whereupon fresh Lenti-X 293 T cells were thawed.

### Cell culture for HEK 293T cells

HEK 293T cells (UCSF Cell Culture Facility) were cultured in medium consisting of Dulbecco’s Modified Eagle Medium (DMEM) (Gibco #10569-010) and 10% fetal bovine serum (FBS) (UCSF Cell Culture Facility). HEK 293T cells were cultured in T75 flasks (Corning #430641U) and passaged upon reaching 80% confluency.

### Cell culture for 3T3 cells

3T3 cells were cultured in medium consisting of Dulbecco’s Modified Eagle Medium (DMEM) (Gibco #10569-010) and 10% fetal bovine serum (FBS) (UCSF Cell Culture Facility). 3T3 cells were passaged upon reaching 80% confluency. To passage, cells were treated with TrypLE express at 37 C for 3 minutes. Then, 10 mL of media was added to quench the reaction and cells were collected into a 50 mL conical tube and pelleted by centrifugation (400xg for 4 minutes). Pellet was resuspended in 5 mL and 1 mL of resuspended pellet was added to a T25 flask (Corning #430639) containing 10 mL of media.

### Culture of mouse embryonic stem cells (mESCs)

mESCs were cultured in “Serum Free ES” (SFES) media supplemented with 2i. SFES media consists of 500 mL DMEM/F12 (Gibco #11320-033), 500 mL Neurobasal (Gibco #21103-049), 5 mL N2 Supplement (Gibco #17502-048), 10 mL B27 with retinoic acid (gibco #17504-044), 6.66 mL 7.5% BSA (Gibco #15260-037), 10 mL 100x GlutaMax (Gibco #35050-061), and 10 mL 100x Pen/Strep. To make “2i SFES”, 1 nM PD03259010 (Selleckchem #S1036), 3 nM CHIR99021 (Selleckchem #S2924) and 1000 units/mL LIF (ESGRO #ESG1106) were added to 45 mL SFES. Prior to use, 1-thioglycerol (MTG; Sigma M6145) was diluted 1.26% in SFES and added 1:1000 to 2i SFES media. To passage, mESCs were treated with 1 mL of accutase in a 6 well plate (Corning #353046) for 5 minutes at room temperature (RT). After incubation, cells were mixed by pipette and moved to a 15 mL conical tube, supplemented with 10 mL SFES and spun at 300xg for 3 minutes. Then, media was removed and cells were counted using the Countess II Cell Counter (ThermoFisher) according to the manufacturer’s instructions. Cells were then plated in 6 well plates that had gelatinized with 1% gelatin for 30 minutes at 37 C at 5 x 10^5 cells per well in 2 mL of 2i SFES. Media was changed every day and cells were split every other day. Cells and culturing reagents were gifted by Dr. Abigail Buchwalter.

### Primary Human T Cell Isolation and Culture

After thawing, T cells were cultured in human T cell medium (hTCM) consisting of X-VIVO 15 (Lonza #04-418Q), 5% Human AB serum and 10 mM neutralized N-acetyl L-Cysteine (Sigma-Aldrich #A9165) supplemented with 30 units/mL IL-2 (NCI BRB Preclinical Repository) for all experiments.

### Lentiviral transduction of primary T cells using LentiX concentrator

Pantropic VSV-G pseudotyped lentivirus was produced via transfection of Lenti-X 293T cells with a modified pHR’SIN:CSW transgene expression vector and the viral packaging plasmids pCMVdR8.91 and pMD2.G using Fugene HD (Promega #E2312). Primary T cells were thawed the same day, and after 24 hr in culture, were stimulated with Dynabeads Human T-Activator CD3/CD28 (Thermo Scientific #11131D) at a 1:3 cell:bead ratio. At 48 hr, viral supernatant was harvested and concentrated using the Lenti-X concentrator (Takara, #631231) according to the manufacturer’s instructions. Briefly, viral supernatant was harvested and potential contaminants were filtered using a 0.45 µM filter (Millipore Sigma #SLHV033RS). Lenti-X concentrator solution was added at a 1:3 viral supernatant:concentrator ratio, mixed by inversion, and incubated at 4 C for at least 2 hours. Supernatant-concentrator mix was pelleted by centrifugation at 1500xg at 4 C for 45 minutes, supernatant was removed and pellet was resuspended using 100 µL media or PBS (UCSF Cell Culture Facility) for each well of T cells. Typically, 2 wells of a 6 well plate was concentrated for 1 well of a 24 well plate plated with 1 million T cells on day of transfection. The primary T cells were exposed to the virus for 24 hr and viral supernatant was exchanged for fresh hTCM supplemented with IL-2 as described above. At day 5 post T cell stimulation, Dynabeads were removed and the T cells expanded until day 12-14 when they were rested for use in assays. For co-culture assays, T cells were sorted using a Sony SH-800 cell or BD FACs Aria sorter on day 5-6 post stimulation.

### Flow cytometry

All flow cytometry data was obtained using a LSR Fortessa or LSRII (BD Biosciences). All assays were run in a 96-well round bottom plate (Fisher Scientific #08-772-2C). Samples were prepared by pelleting cells in the plate using centrifugation at 400xg for 4 minutes. Supernatant was then removed and 200 µL of PBS (UCSF Cell Culture facility) was used to wash cells. The cells were again pelleted as described above and supernatant was removed. Cells were resuspended in 120 µL of Flow buffer (PBS + 2% FBS) and mixed by pipetting prior to flow cytometry assay.

### Inhibitor Assays

100,000 cells were plated in a 96 well round bottom plate with either 5 µM MG132 (Sigma-Aldrich #M7449-200UL), 1 µM MLN4924(Active Biochem #A-1139), 100 nM Bafilomycin A1(Enzo Life Sciences #BML-CM110-0100), or DMSO vehicle control and incubated at 37 C for 5 hours. After incubation, cells were pelleted by centrifugation at 400xg for 4 minutes. Supernatant was then removed and cells were washed with 200 µL PBS. Cells were pelleted again (400xg for 4 minutes) and resuspended in flow buffer (PBS + 2% FBS) for assay by flow cytometry.

### Antibody staining

All experiments using antibody staining were performed in 96 well round bottom plates. Cells for these assays were pelleted by centrifugation (400xg for 4 minutes) and supernatant was removed. Cells were washed once with 200 µL of PBS and pelleted again by centrifugation (400xg for 4 minutes) and the supernatant was removed. Cells were resuspended in a staining solution of 50 µL PBS containing fluorescent antibody stains of interest. Anti-myc antibodies (Cell Signaling Technologies #2233S, #3739S and #2279S) was used at a 1:100 ratio while antiV5 (ThermoFisher Scientific #12-679642) and antiFLAG (R&D Systems #IC8529G-100) antibodies were used at a 1:50 ratio for flow cytometry assays. For FACS, all antibodies were used in a 1:50 ratio in 100 uL.

### Intracellular staining

For intracellular staining assays, cells were pelleted at 400xg for 4 minutes following co-culture. After careful removal of supernatant, cells were treated with Zombie UV fixable Viability kit diluted 1:500 in PBS for 30 minutes in the dark at room temperature. Cells were then fixed and permeabilized using the eBioscience Foxp3/Transcription Factor Staining Buffer Set (ThermoFisher #00-5523-00) following manufacturer’s recommended procedure. Following the fixation/permeabilization process, cells were stained with the ZAP-70 (136F12) Rabbit mAb (PE Conjugate) (Cell Signaling #93339) antibody for 1 hour at room temperature in the dark. The stain was then washed off with the perm/wash buffer in the eBiosciences kit twice and resuspended in flow buffer. Cells were then analyzed by flow cytometry.

### CRISPR/Cas9 HDR Knock-in template generation

CRISPR/Cas9 knock-in template was generated using PCR from a plasmid encoding the homology arms and desired knock-in payload as described in Shy et al. Double-stranded DNA was purified by solid phase reversible immobilization (SPRI) bead cleanup using AMPure XP beads (Beckman Coulter #A63881). Beads were added in 1.8:1 volume:volume ratio to PCR template and isolated per manufacturer’s instructions. For GFP-ZAP70 template, HDR template was eluted in 30 µL of water.

### CRISPR/Cas9 RNP formulation

RNPs were produced following protocols described in Shy et al. crRNA and tracRNA were synthesized by IDT and resuspended in provided buffer at 160 µM and kept as 3 µL aliquots at −80 C. All incbuation steps described in this section were performed on a heatblock. gRNA was made by mixing crRNA and tracrRNA at 1:1 v/v ratio and annealed by incubation for 20 minutes at 37 C. ssDNAenh (IDT) was added at 0.8:1 v/v ratio and mixed by pipette. The ssDNAenh electroporation enhancer is a sequence described by Shy et al. shown to improve knock-in efficiency. The ssDNAenh (5’-TCATGTGGTCGGGGTAGCGGCTGAAGCACTGCACGCCGTACGTCAGGGTGGTCACGAGGGTGGGCCAGGGCACGGGCAGCTTGCCGGTGGTGCAGATGAACTTCAGGGTCAGCTTGCCGTAGGTGGC-3’) was synthesized by IDT, resuspended to 100 µM in duplex buffer and stored at −80 °C in 5 µL aliquots. Then, 40 µM of Cas9-NLS (Berkeley QB3 Macrolab) was added to the gRNA+ssDNAenh mixture at a 1:1 v/v ratio. This results in a final molar ratio of sgRNA:Cas9 of 2:1. This mixture was mixed by pipette and incubated for 15 minutes at 37 C. Based on a Cas9 protein basis, 50 pmol of RNP was used for each electroporation.

### Electroporation

Electroporation was done 7 days after stimulation by DynaBeads using a P3 Primary Cell 4D-Nucleofector™ X Kit (Lonza #V4XP-3012). 750 ng of HDR template was mixed with 50 pmol of RNP and incubated at room temperature for 10 minutes. During this incubation, DynaBeads were removed from cells and cells were spun at 200xg for 7 minutes. Supernatant was removed and cells were resuspended in 20 µL of Lonza P3 buffer. RNP+HDR template mixture was added to cells and 20 µL of this mixture was then added to the 96-well electroporation vessel. Cells were then electroporated using a Lonza 4D-Nucleofector^®^ X Unit (Lonza #AAF-1003X) with code EH-115. Immediately, 90 µL of warm hTCM+IL-2 was added to cells and cells were then incubated for 20 minutes at 37 C. Cells were then transferred to a fresh 96 well plate and diluted to 1.0 × 10^6^ cells per ml in hTCM+IL-2 and 0.05 µM of the small molecule inhibitor TSA (Cayman Chemical). The 96 well plate was then spun at 200xg for 7 min and incubated in a tissue culture incubator overnight. 24 hours following TSA treatment, TSA-containing media was removed and fresh hTCM+IL-2 was added. Fresh cytokines and media were added every 2-3 days until sorting by FACS.

### Co-culture assays

For all assays, T cells and target cells were co-cultured at a specified effector to target ratios with cell numbers varying per assay. All assays contained between 10,000 and 50,000 of each cell type. The Countess II Cell Counter (ThermoFisher) was used to determine cell counts for all assays set up. T cells and target cells were mixed in 96-well round bottom tissue culture plates in 200 µL T cell media, and then plates were centrifuged for 1 min at 400x g to initiate interaction of the cells prior to incubation at 37 C.

### Incucyte cell lysis assays

Primary human CD8+ T cells and target cells were plated in a flat bottom 96 well plates (cat num.). For suspension target cells, wells were coated with 50 µL of 5 µg/mL fibronection and incubated for 1 hour at room temperature. Fibronectin solution was then removed and plates were left to dry in a biosafety cabinet for 1 hour. Engineered T cells and target cells are then added at effector:target ratios as needed and allowed to settle at room temperature for 30 minutes. Images were taken every 3 hours using Incucyte hardware and software over the course of the experiments.

### Data analysis

Data analysis was performed using the FlowJo software (FlowJo LLC.) and Python. For co-culture assays, desired cell populations were isolated by FACS using a Sony SH800 or Aria (BD) cell sorter. For non co-culture assays, desired cell populations were isolated by gating in FlowJo following flow cytometry. Incucyte data was analyzed and quantified by Incucyte software and plotted using Python.

## Acknowledgements

The authors thank members of the El-Samad Lab for helpful comments and discussion regarding the manuscript. The authors would also like to thank the Wendell Lim Lab and the Cell Design Institute for providing reagents and support for primary T cell experiments. We thank Brian Shy, Alvin Ha and the Alex Marson Lab for their guidance and help with reagents on CRISPR/Cas9 experiments and Ray Liu, Iowis Zhu and the Kole Roybal Lab for reagents and advice on the SNIPR tool. We also thank Drs. Itay Koren and Stephen Elledge for providing sequences and advice for experiments using dominant negative cullins. We thank Harold Marin and the Abigail Buchwalter lab for their help and advice with mESC work. This worked was supported by the Defense Advanced Research Projects Agency (DARPA) (HR0011-16-2-0045), NIH/NCI/NIBIB (U54CA244438), NIH/NCI (U01CA265697) and the Chan-Zuckerburg Initiative.

## Author Contributions

M.S.K., W.A.L., H.E.-S., and A.H.N. conceived and designed the study and experiments. M.S.K. and A.H.N. designed sequences and vectors. M.S.K., H.K.B., G.E.S., and A.H.N. performed and analyzed experiments. W.A.L. and H.E.-S. provided funding support. M.S.K., H.E.-S., and A.H.N. wrote the manuscript.

