## Supplemental Figures for "Degron-based bioPROTACs for controlling signaling in CAR T cells"

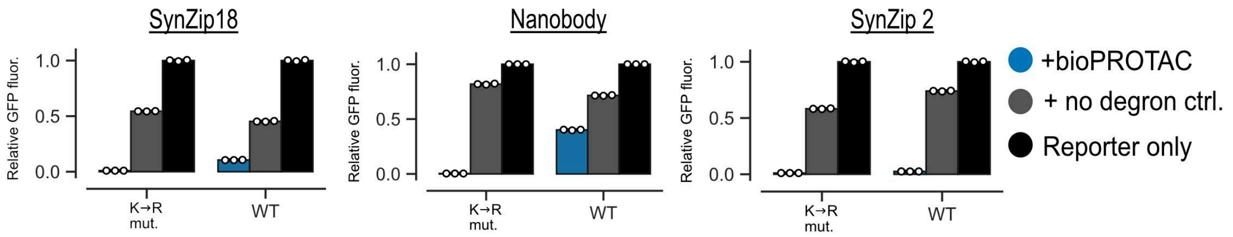


Figure S1. **bioPROTA**C **degradation improves upon mutation of lysine residues to arginine.** Mutation of the lysine residues to arginine in the binding domain of bioPROTACs improves degradation of GFP. GFP fluorescence was measured by flow cytometry. Relative GFP fluorescence is calculated as described for Fig 1. Labels above plots describe the binding domain used by each bioPROTAC. Dots represents technical replicates and error bars shows SEM. Data is representative of three independent experiments.

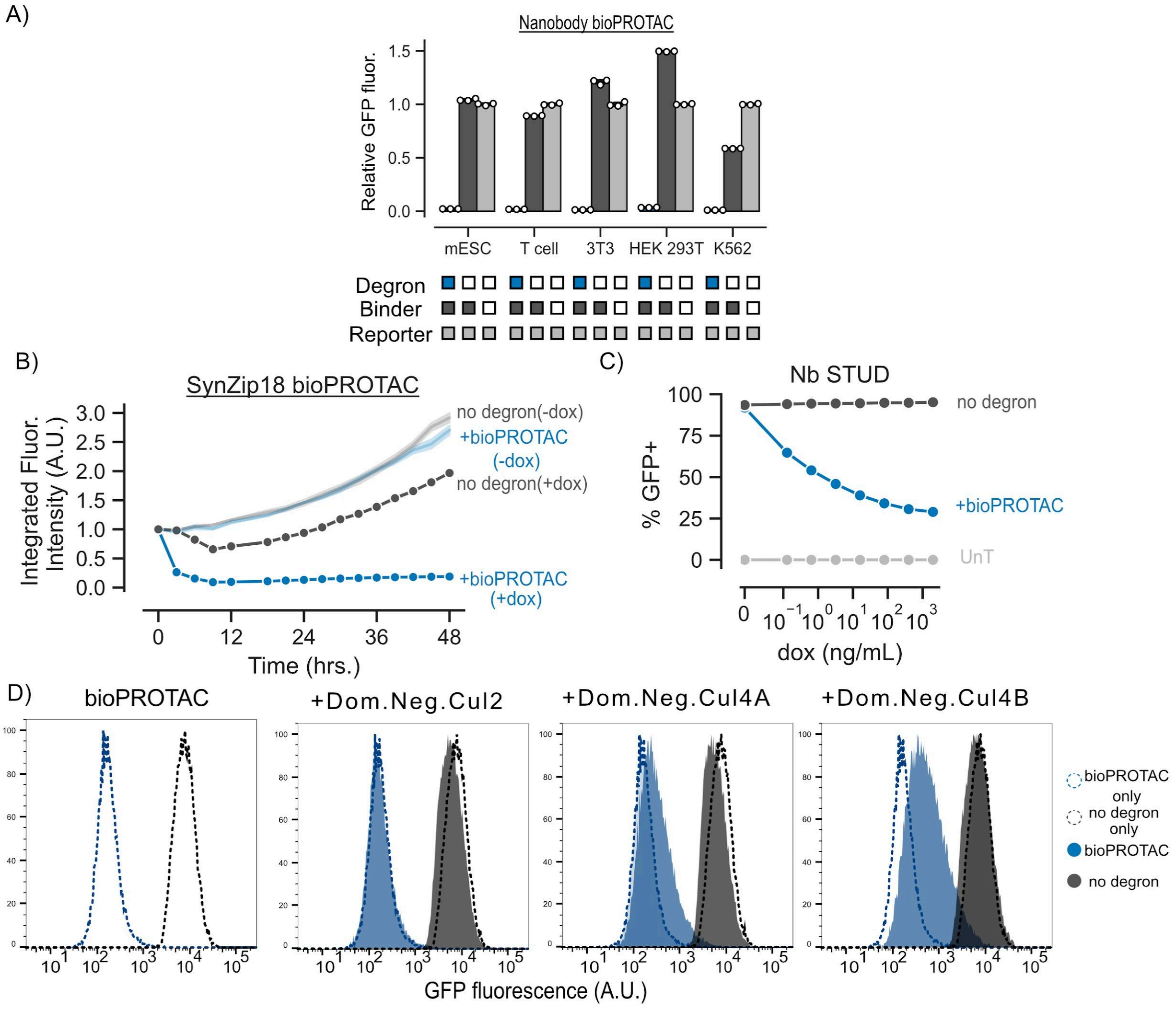

Figure S2. **Characterization of bioPROTAC mechanism and function.** (A) The vhhGFP4 nanobody bioPROTAC showed potent degradation across all tested mammalian cell lines. GFP fluorescence was measured by flow cytometry. Relative GFP fluorescence is calculated as described in Fig 1. Dots represents technical replicates and error bars show SEM.
(B) Time traces of SynZip bioPROTAC degradation with additional controls. GFP fluorescence measurements were captured by fluorescence microscopy and live cell imaging on an Incucyte machine. Data was analyzed in the Incucyte software. Data is normalized to the time 0 values for each condition. (C) The vhhGFP4 Nanobody bioPROTAC showed dose-dependent degradation. bioPROTACs titration results in dose-dependent degradation of cytosolic proteins. Jurkat T cells were lentivirally transduced with a GFP reporter protein and a plasmid encoding a doxycycline inducible nanobody bioPROTAC and the Tet3G protein. After isolation by FACS, cells were treated with a 2-fold titration series of dox starting at 2000 ng/mL or a media only control for 48 hours. bioPROTAC efficacy was assessed by flow cytometry. Each dot is the mean of three biological replicates. Error shows SEM. (D) Dominant negative cullin ring ligases from the Cul4 family rescued bioPROTAC degradation. GFP fluorescence was measured by flow cytometry. Plots are representative of three replicates.

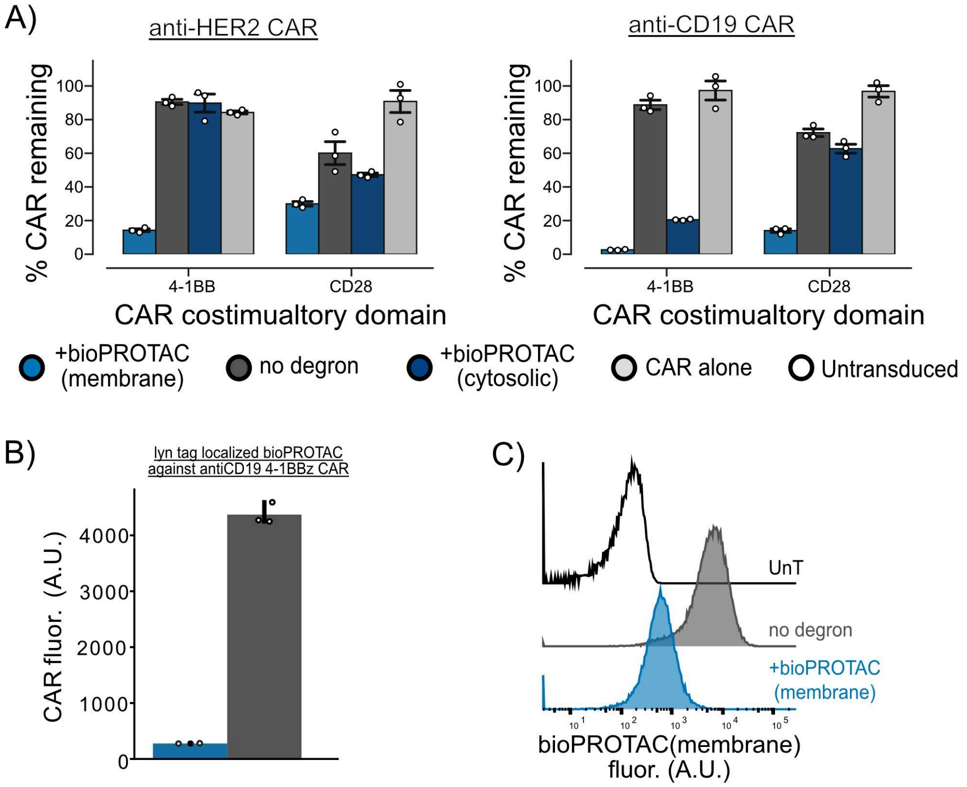


Figure S3. **bioPROTAC(membrane) characterization.** (A) bioPROTAC(membrane) degradation performs better against second-generation CARs with 4-1BB than those with CD28 costimulatory domains. Jurkat T cells lentivirally were transduced with plasmids encoding a second-generation CAR and a bioPROTAC(membrane) as described in Figure 3A. CAR expression was measured by staining for an extracellular tag fused to the CAR and measured by flow cytometry. (B) bioPROTAC(membrane) can be recruited to the membrane with different domains without loss of function. Jurkat T cells were engineered to express an antiCD19 4-1BB CAR. Then, either a lyn tagged bioPROTAC(membrane), a lyn tagged no degron control or the DAP10ss bioPROTAC(membrane) described in Figure 3 was introduced by lentivirus. CAR expression levels were assessed by antibody stain for an extracellular tag. (C) Flow cytometry histograms representing for DAP10 bioPROTAC(membrane) expression in Jurkat T cells. bioPROTAC(membrane) fluorescence value were measured by immunofluorescence staining for a V5 tag fused to the extracellular domain of the protein followed by flow cytometry.

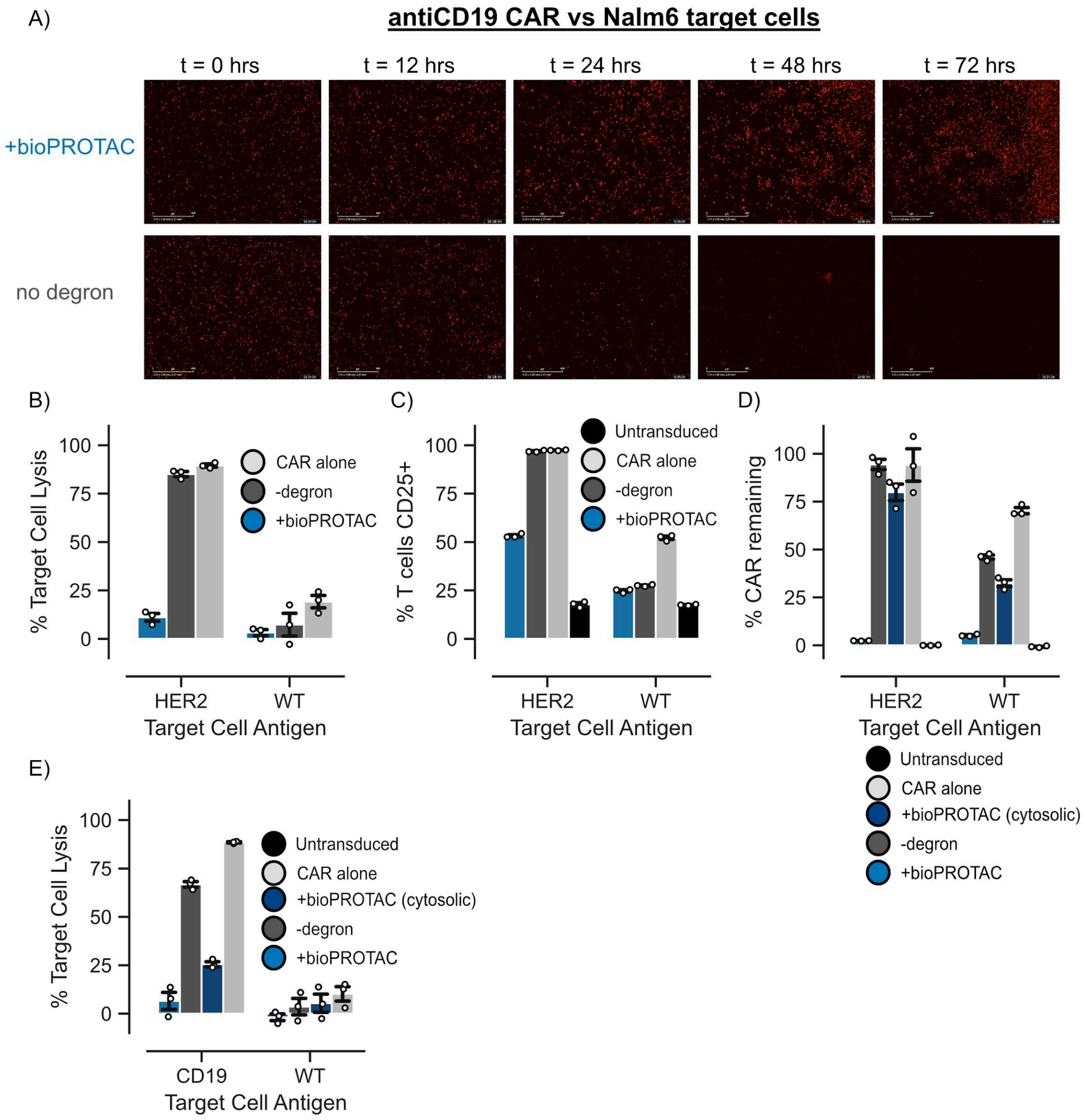

Figure S4. **Further characterization of bioPROTAC(membrane) control of CARs.** (A) Representative images of live cell imaging of target cell lysis by CAR T cells cocultured with mCherry+ Nalm6 cells. Images are taken by an Incucyte machine. (B) bioPROTAC(membrane) impairs antiHER2 4-1BBz CAR cytotoxicity. Cell lysis was calculated using cell counts measured by flow cytometry and normalized to cell counts of UnT cells against each target cell type. Dots represent technical replicates and error bars show SEM. (C) bioPROTAC(membrane) prevents antiHER2 4-1BBz CAR associated activation of CD25. CD25 levels were measured by immunostaining followed by flow cytometry. Dots represent technical replicates and error bars show SEM. (D) bioPROTAC(membrane) induces internalization of antiHER2 4-1BBz CAR. CAR fluorescence was measured by immunostaining following by flow cytometry. Dots represent technical replicates and error bars show SEM. (E) Membrane localization is required for strong ablation of CAR induced cytotoxicity in primary T cells. Same data as Figure 4A including a control with a soluble bioPROTAC.

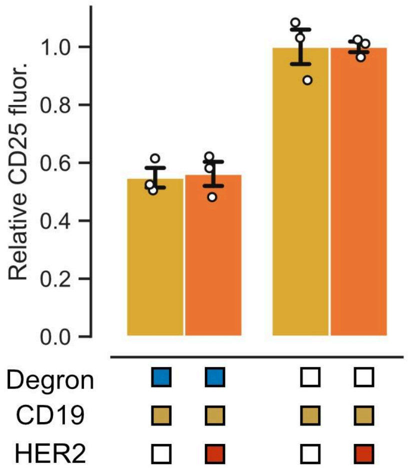


Figure S5. **bioPROTAC circuit does not affect CD25 levels following the first challenge.** CD25 levels of bioPROTAC expressing were assessed by antibody stain after 48 hours of co-culture with CD19 or CD19 and HER2 expressing K562 cells . CD25 fluorescence levels were measured by flow cytometry. Relative CD25 levels were calculated by normalizing each test condition to a no degron control. Each dot represents a technical replicate. Error bars show SEM.

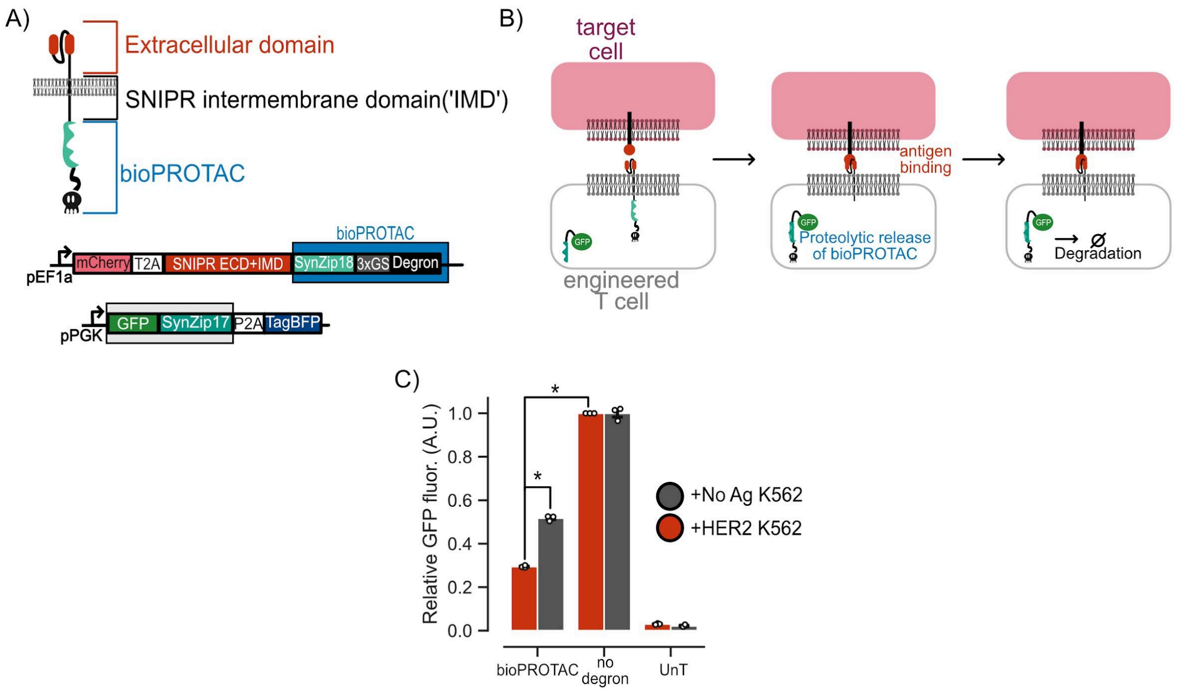


Figure S6. **bioPROTACs are synergistic with existing protein tools.** (A) Schematic of domain swap of transcriptional activation domain of SNIPR to SynZip18 bioPROTAC to create antigen inducible degradation tool. CD4+ primary human T cells were transduced with the diagrammed lentiviral payloads. Engineered T cells were then co-cultured with HER2 expressing K562s or no antigen K562s for 72 hours. SNIPR-bioPROTAC fusion protein activity was measured by changes in GFP fluorescence by flow cytometry. (B) Cartoon depicting proposed mechanism of SNIPR-bioPROTAC fusion protein. (C) Combining bioPROTACs with the extracellular and intermembrane domains of SNIPR creates an antigen inducible targeted protein degradation tool. GFP fluorescence was measured by flow cytometry. Relative GFP fluorescence was calculated by normalizing each test condition to a reporter only control. Each dot represents a technical replicate and error bars show SEM. Significance determined by paired t-test, * = P < .01
